# Unsupervised neuronal cell typing from connectivity by minimum description length

**DOI:** 10.64898/2026.10.20.760604

**Authors:** Gregory Schwartzman

## Abstract

Connectomes are being reconstructed at rapidly increasing scale, creating a growing need for automated methods to identify neuronal cell types. Existing approaches rely heavily on expert annotation or on reference connectomes, and reliable unsupervised typing remains an open problem. We present a fully unsupervised method that recovers neuronal types using only connectivity, achieving 93–98% agreement with expert annotations on fruit fly optic lobes and visual systems. This is the first result to show that connectivity alone can achieve *near perfect* recovery of neuronal types in large regions of the brain.

Our method is based on the *minimum description length* (MDL) principle: a good partition of neurons into clusters corresponds to a *compression* of the connectome. This provides a principled theoretical foundation for neuronal typing through the lens of compression. It also opens a path toward unsupervised typing in substantially harder regions, including the central brain and ventral nerve cord.

## Introduction

Connectome reconstruction maps neurons and their synaptic connections in the brain. This is typically accomplished using electron microscopy (EM) imaging, automated segmentation and synapse detection, followed by human proofreading and annotations of *cell types*. A neuronal cell type is a population of neurons that share similar molecular, anatomical and physiological properties. Cell types provide a basis for interpreting neuronal circuits and comparing their organization across species. By organizing large connectomes into cell types, they allow circuit structure to be studied at a scale beyond individual neurons. Connectome reconstructions now span the adult *Drosophila melanogaster* (fruit fly, a popular model organism) brain [1, 2], visual system [3, 4], and ventral nerve cord [5], as well as unified brain and ventral nerve cord connectomes of male [6] and female flies [7]. Beyond insects, synaptic connectivity has also been reconstructed in the zebra finch basal ganglia [8] and mouse visual cortex [9]. With the recent rise of large-scale connectomics datasets, there is a growing need for *fully-automated* methods to identify neuronal cell types.

### Cell typing

Current approaches to cell typing depend on whether an annotated reference of the same species and brain region is available. When such a reference exists, it provides a catalogue of types for the new dataset. Neurons can be assigned to these types through morphological comparison, commonly using NBLAST [10], together with connectivity analysis and expert review [2]. Recently it was shown [11] that connectivity and a small subset of neurons labeled by type suffice to recover the remaining neurons’ types in the fly optic lobes with near-perfect accuracy. Connectivity-based alignment also enables cell-type transfer between male and female ventral nerve cords and identification of sex differences between them [12].

Without a suitable reference—for example, in a new species or a previously untyped brain region—the types must be discovered from the data. This is a *graph clustering problem*: the vertices are the neurons, the edges their connections, and the clusters the types. The established practice is morphological clustering refined by connectivity and extensive expert curation, which took large teams years for the fly brain [2–4]; for the larval fly brain, where morphology is less informative, types were discovered by clustering connectivity directly [13] followed by expert validation. The state of the art for unsupervised typing is the NTAC algorithm [11]. While it is able to reconstruct types that contain many neurons reasonably accurately, it struggles on smaller types.

### Overview of results

In this work, we present MOMONGA (MDL Optimization for Mapping Of Neuronal Groups from Adjacency), a fully unsupervised method for discovering neuronal cell types from connectivity alone. MOMONGA uses the minimum description length principle [14, 15] to identify types through the compression afforded by their shared connectivity. Roughly speaking, for our setting the MDL principle states that the best partition of neurons into clusters is the one that allows the connectome to be described most concisely. This provides a single criterion for selecting both the partition and its number of clusters, without requiring any labeled neurons.

Evaluated on state-of-the-art female and male fruit fly connectomes, MOMONGA achieves 93–98% accuracy in the optic lobes and visual systems, against 69–88% for NTAC, without requiring any labeled neurons. The visual system holds about two thirds of the neurons of the fly brain but only about a tenth of its cell types (Supplementary Table 1); most types are in the central brain and nerve cord, where each has only a few neurons. Although MOMONGA considerably outperforms NTAC in the central brain (72% vs 54% accuracy) and ventral nerve cord (54% vs 45%), these regions still remain challenging. To alleviate this issue, MOMONGA also provides a confidence score for each neuron. In the central brain, retaining the top 25% of neurons by confidence yields a clustering with pure representatives of more than 50% of all expert types. Less than 5% of the retained clusters are impure or duplicates. This partial catalogue could provide a starting point for accelerating cell typing in new connectomes. Accuracy, macro-F1, cluster purity and duplicates are defined at the start of Results. Accuracy results and confidence-filtered catalogues are shown in Figs. 1 and 2, respectively.

**Figure 1.**
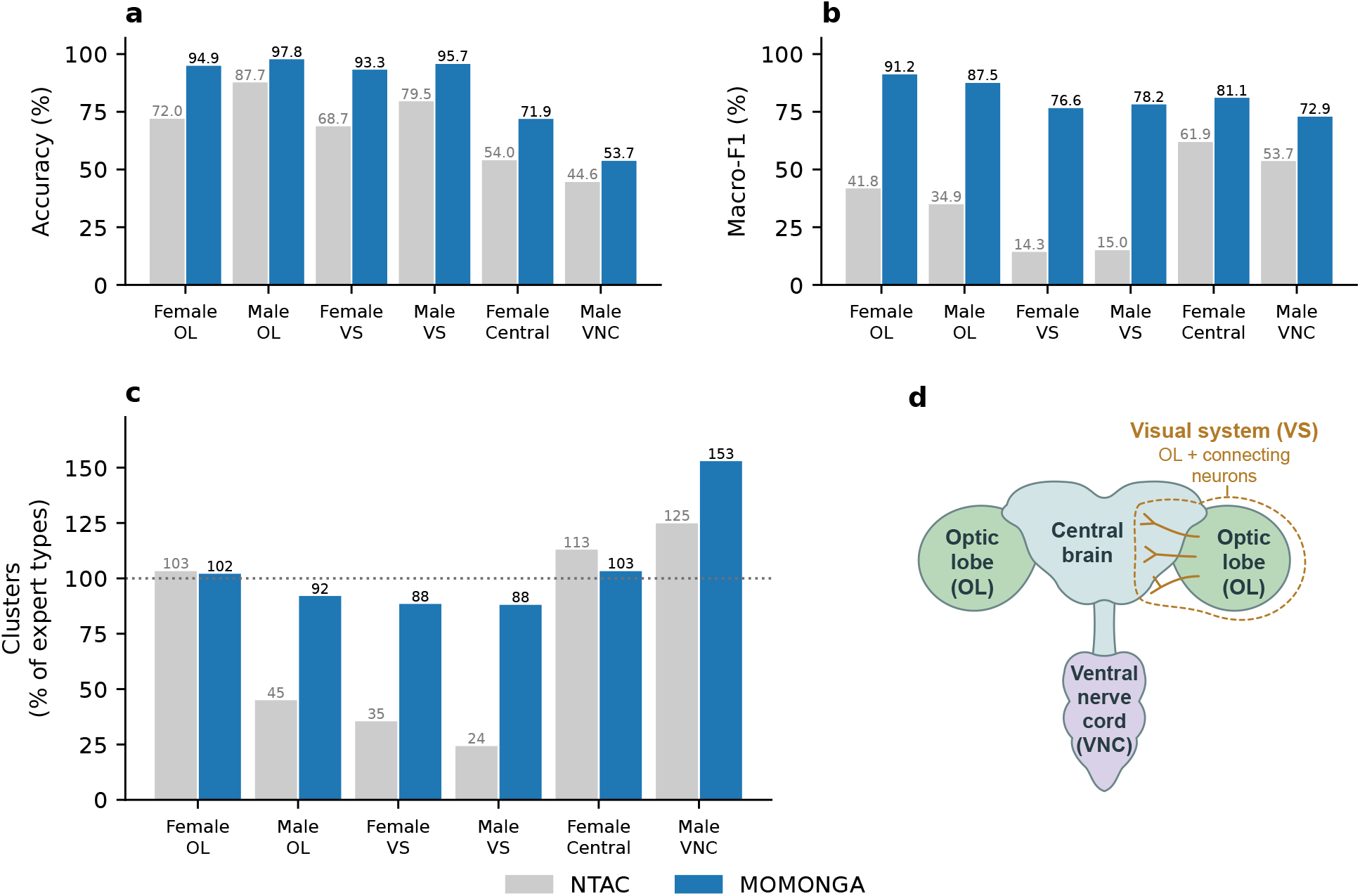
Cell-type recovery by NTAC and MOMONGA: accuracy (a), macro-F1 (b), and number of clusters the method returns as a percentage of total expert types (c). Visual results are evaluated separately for each hemisphere and averaged with equal weight. Female Central and Male VNC results do not separate hemispheres. The schematic (d) identifies the anatomical regions; the visual system includes the optic lobe and connecting neurons. All datasets omit connections with fewer than 5 synapses in either direction. Accuracy and macro-F1 are defined in the Accuracy metrics paragraph in Results.

**Figure 2.**
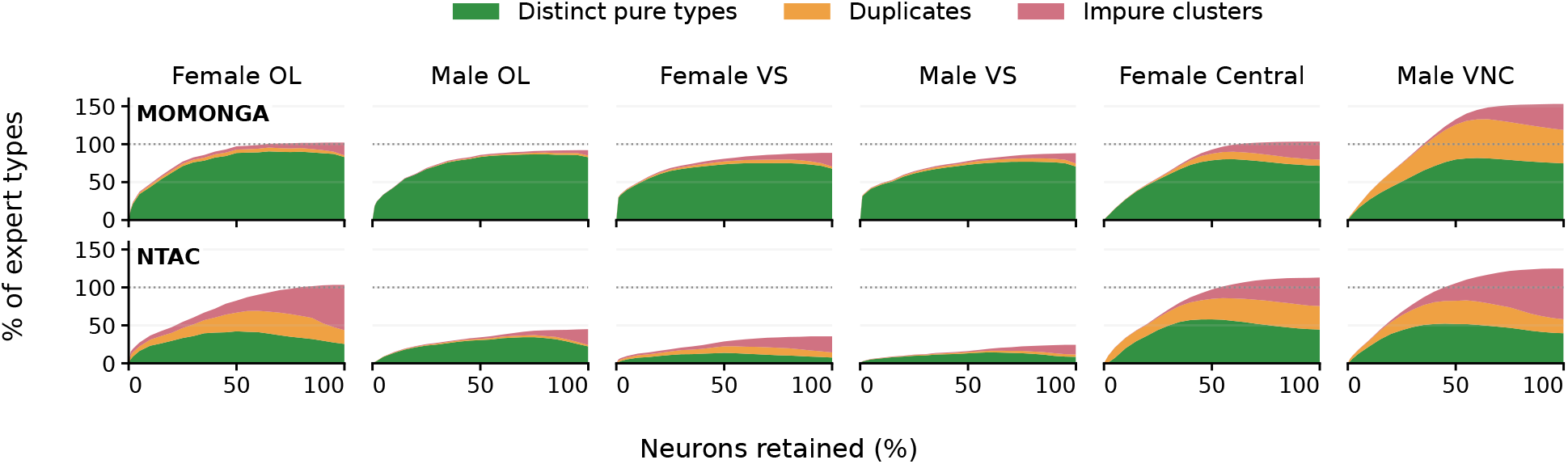
Cell-type catalogues produced by MOMONGA and NTAC after confidence filtering. The plots show distinct pure types, duplicate pure clusters and impure clusters as percentages of the total number of expert types, against the percentage of neurons retained. Neurons are retained in decreasing order of confidence, defined in Appendix A.1; purity is measured among the retained neurons. Visual results are evaluated separately for each hemisphere and averaged with equal weight. Female Central and Male VNC results do not separate hemispheres. Purity and duplicates are defined in the Accuracy metrics paragraph in Results.

## Results

### Accuracy metrics

Because unsupervised methods return clusters without type labels, evaluating their accuracy requires matching clusters to expert types. We use a one-to-one matching that maximizes the number of correctly assigned neurons, computed with the Hungarian algorithm [16]. Accuracy is the fraction of neurons assigned to their correct expert type after this matching; neurons in unmatched clusters count as errors. Using the same matching, macro-F1 is the average F1 score across all expert types, giving each type equal weight regardless of its size. Unmatched expert types receive an F1 score of zero. We also measure cluster purity: the fraction of neurons belonging to the most common expert type in a cluster. A cluster is pure if this fraction is at least 95%, and impure otherwise. When several pure clusters have the same majority expert type, all but one are counted as duplicates. The total number of duplicates is summed over expert types.

### The challenge

Connectivity-based typing rests on the observation that neurons of the same type have similar *connectivity profiles* : patterns of incoming and outgoing connections to other types [3, 4, 11]. These profiles act as a fingerprint for the type (Fig. 3). Without known types, however, these profiles cannot be computed directly. A proposed partition determines which connections are combined into each profile, so changing the partition also changes the profiles being compared. Unlike clustering with fixed feature vectors, both the clusters and the profiles used to compare their members must therefore be inferred to-gether.

**Figure 3.**
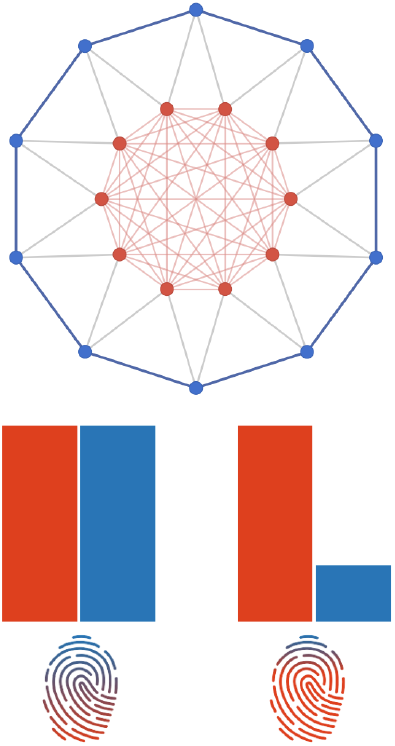
A simplified undirected graph with two neuronal types, red and blue. Blue neurons connect equally to both types, whereas red neurons connect mainly to red neurons. The bars show the connectivity profiles of the blue type (left) and red type (right), which act as fingerprints for the two types.

NTAC heuristically optimizes an objective function that measures the total disagreement between each neuron’s connectivity profile and the representative profile (mean or median) of its assigned cluster [11]. This objective does not directly penalize the number of clusters— putting every neuron in its own cluster gives zero cost. Furthermore, NTAC constructs partitions by greedily adding one cluster at a time and reassigning neurons, without merging existing clusters. Although it can select an earlier partition along this sequence, it does not refine a partition by both increasing and decreasing its number of clusters. We therefore seek an objective that balances similarity within clusters against the cost of additional clusters, together with an algorithm that allows both splitting and merging of clusters.

### Our approach

We use the minimum description length (MDL) principle [14, 15] to define this balance and design our algorithm. To illustrate this principle, suppose Alice wants to send Bob the graph in Fig. 3. One option is to send the vertices and then list every edge separately. Another is to send the vertices, their division into red and blue groups, and the connectivity profile shared by each group: every blue vertex connects to two red and two blue vertices, while every red vertex connects to the other red vertices and two blue vertices. She then specifies which particular vertices form these connections. The group information helps when the resulting savings outweigh the cost of describing the groups.

The MDL principle formalizes Occam’s razor by favoring explanations that give the shortest combined description of the explanation and the data. In our context, the data are the connectome, and the explanation is a partition of neurons into clusters. We therefore seek a partition that allows the connectome to be described concisely, including the description of the types themselves. To make this criterion concrete, we specify an explicit encoding of the graph and its partition, whose length can be evaluated exactly for any proposed partition. Our encoding is not the shortest possible, and it could be optimized further, but a shorter code does not necessarily give better typing: what matters is not the absolute length but how the code ranks competing partitions, for example whether merging two clusters shortens the description. We use a heuristic search to find partitions with shorter descriptions under this encoding. This provides a single criterion for selecting both the partition and its number of clusters, without requiring any labeled neurons.

### Overview of MOMONGA

MOMONGA is an unsupervised method for identifying neuronal cell types from connectivity alone. It applies the MDL principle through an explicit code describing the connectome and a partition of its neurons. Formally, given the connectome as a directed weighted graph, we seek a partition *P* of the connectome, into any number of clusters, that minimizes *L*(*P* ), the total description length under this code. The length can be evaluated exactly for any proposed partition and consists of four terms:

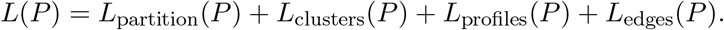

Here, *L*_partition_(*P* ) describes which neurons belong to each cluster; *L*_clusters_(*P* ) describes the total connection weights between clusters; *L*_profiles_(*P* ) describes each neuron’s connection weights to and from the clusters; and *L*_edges_(*P* ) specifies which individual neurons are connected and the weights of their connections. Specifically, *L*_partition_(*P* ) and *L*_clusters_(*P* ) account for the cost of describing the types and their aggregate connectivity, while *L*_profiles_(*P* ) rewards similarity between the profiles of neurons within each cluster. Each stage uses the information supplied by the preceding stages (Fig. 4). The full encoding is described in Methods.

**Figure 4.**
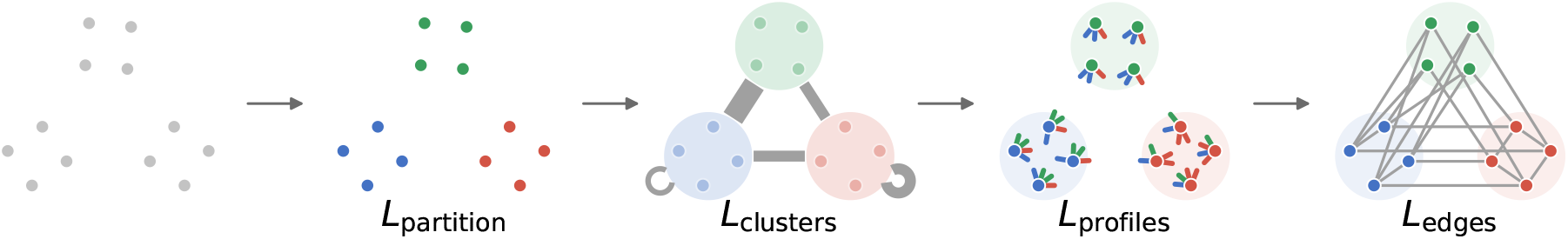
An intuitive illustration of the code, showing what the receiver knows after each stage of the code, on a simple undirected and unweighted graph with three clusters. At the start it knows the vertices. Each stage then adds one piece of information: the partition; the number of edges between each pair of clusters; the number of edges from each vertex to each cluster; and finally the individual edges. Below each panel is the term of *L*(*P* ) that pays for that stage.

MOMONGA has two functions: refinement and transfer. Refinement takes a coarse initial partition and iteratively improves it by alternating between split and merge rounds. In a split round, we heuristically split each cluster into two, aiming to separate members with different connectivity profiles. In a merge round, we iteratively merge pairs of clusters as long as each merge decreases the description length. The algorithm terminates when a merge round does not improve the description length relative to the previous merge round, and returns the previous round’s partition.

Transfer takes two graphs and their partitions as inputs. These correspond to similar anatomical regions, either from different specimens or from opposite sides of the same connectome. We choose the partition with the smaller description length per edge as the donor, using its clusters to guide neuron assignments in the recipient. We first align donor and recipient clusters, aiming to match clusters that correspond to the same cell type. After alignment, we reassign recipient neurons between clusters using the donor as a guide. We consider several alignments and select the resulting recipient partition with the shortest joint description of the two graphs. Finally, we move individual recipient neurons between clusters whenever doing so further decreases this joint description length.

We evaluate MOMONGA on expert-annotated adult fruit fly connectomes: the female brain’s optic lobes, visual systems and central brain [1–3], the male optic lobes and visual systems [4, 6], and the male ventral nerve cord [5].

The full MOMONGA pipeline starts from several coarse partitions generated by NTAC. We refine each starting partition and select the result with the smallest description length as the output. For the optic lobes and visual systems, we refine the left and right sides separately, then apply transfer between the left and right partitions of each region. The central brain and nerve cord are each refined as a single graph containing neurons from both the left and right sides. For these regions, we exploit left–right symmetry by matching neurons across sides and using the resulting correspondence to augment the connectivity profiles used to initialize refinement.

## Related work

### Cell typing from connectivity

Winding et al. [13] used connectivity to embed neurons of the larval fly brain and clustered the embedding hierarchically. Cell typing in adult fly connectomes has generally combined morphological comparison, connectivity analysis and expert curation [2– 4, 17]. In the female visual system, Matsliah et al. [3] iteratively refined type assignments by comparing neurons’ connectivity to types using weighted Jaccard distance. For the male visual system, Nern et al. [4] applied Ward clustering to connectivity profiles, with morphology and genetic lines helping resolve ambiguous assignments.

Unseeded NTAC [11] formulated connectivity-based typing as approximate equitable partitioning. Unlike clustering a fixed connectivity embedding, its profiles change with the partition. The algorithm grows the number of clusters by adding seeds and reassigning neurons to improve a within-cluster similarity objective. MOMONGA uses this algorithm to generate starting partitions, then refines them under a description-length objective. Its split and merge operations allow the number of clusters to increase or decrease, while description length provides a criterion for selecting among the refined partitions.

Connectivity-based transfer between connectomes has previously used individual-neuron alignment [12]. MOMONGA aligns clusters, reducing the size of the matching problem, and uses description length to select and refine the transferred assignments.

### Description length

The minimum description length principle selects an explanation through the combined length of its description and that of the data given the explanation [14, 15]. Wallace and Boulton introduced a related coding principle for clustering [18]. For networks, Rosvall and Bergstrom selected community partitions through the compression they afford [19]. MDL was also highlighted as a promising future direction for neuronal classification once sufficiently large, statistically representative datasets became available [20]. That review did not implement or evaluate an MDL-based typing method.

### Cell-type recovery with MOMONGA

We evaluate MOMONGA on expert-annotated adult fruit fly connectomes against NTAC, the current state of the art for unsupervised cell-typing [11]. In Fig. 5, we compare MOMONGA with NTAC on each dataset as a function of the number of clusters in the NTAC starting partition^1^. We ran NTAC up to 100 clusters beyond the expert-type count for the optic lobes and visual systems, and 1,000 beyond it for the central brain and ventral nerve cord, to test whether overestimating the number of types affects the final output of MOMONGA. For efficiency, we evaluate MOMONGA on a subset of the NTAC trajectory. For each dataset, we select the output partition with the smallest description length. The NTAC baseline is the partition with the smallest Jaccard cost over its full trajectory—the same selection criterion as in [11].

**Figure 5.**
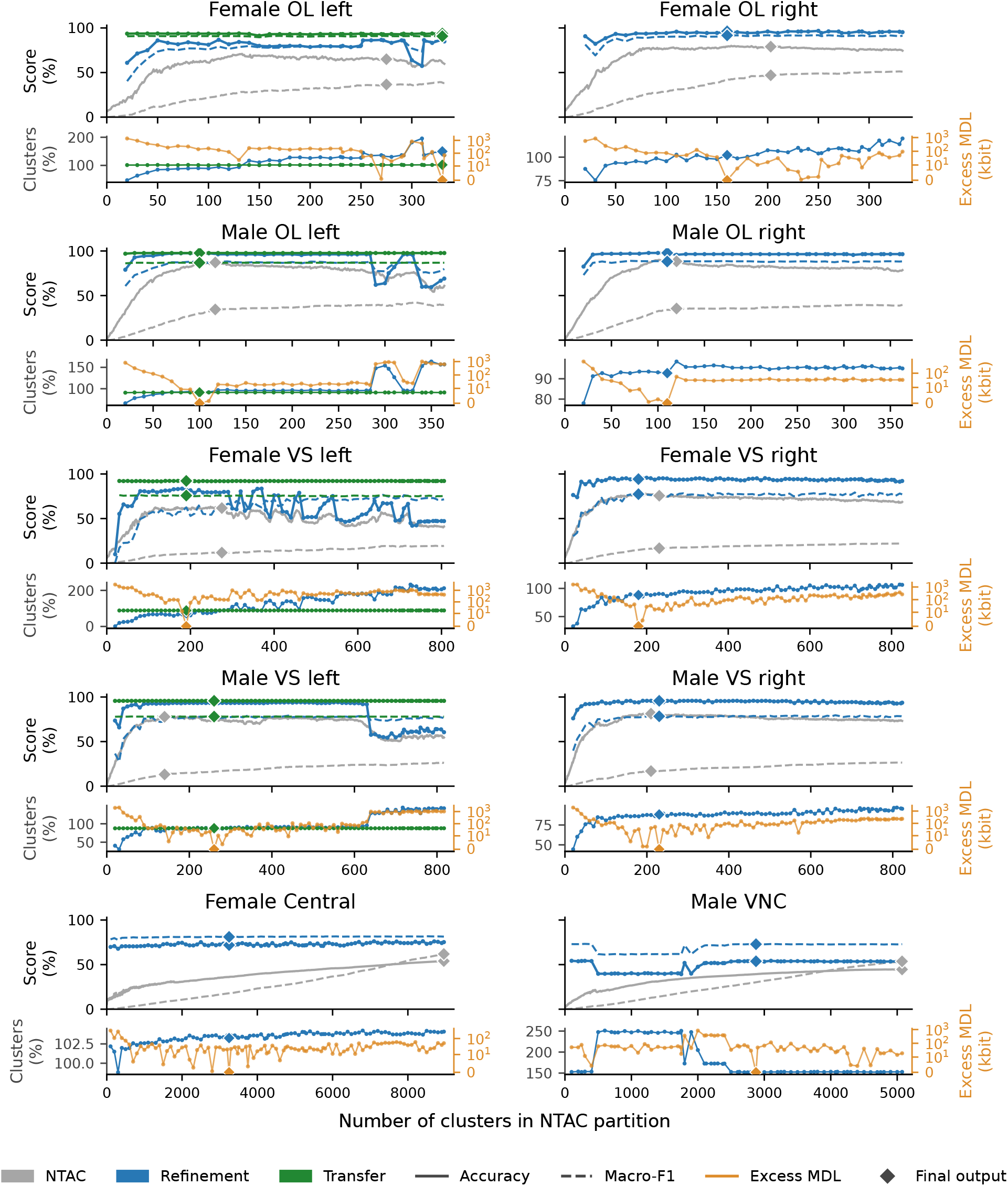
Refinement and transfer across NTAC starting partitions. Upper panels compare accuracy and macro-F1 across the full NTAC trajectory and completed refinements and transfers. Diamonds mark the final outputs: the minimum-Jaccard NTAC partition and the MDL-selected MOMONGA result, after transfer where applicable. Lower panels show the difference between each refined partition’s description length and the minimum across the grid for that dataset. They also show the cluster counts of the refined and transferred partitions as a percentage of expert types. Diamonds in the lower panels mark the description length and cluster count of the MDL-selected refined partition and, where applicable, the cluster count of its transferred partition. Refinements are evaluated every 10 NTAC steps (every 100 for Female Central and Male VNC). NTAC and refinement are deterministic; transfer curves and diamonds are means over five sets of random seeds.

For the optic lobes and visual systems, we additionally apply transfer between hemispheres. Among the refined partitions across both sides, we choose the partition with the smallest description length divided by the number of edges as the donor. Keeping this donor fixed, we apply transfer to each recipient grid point. The algorithm correctly selected the right side as the donor in all pairs. Expert labels were not used in any stage of refinement or transfer, and the final outputs were selected by description length alone.

MOMONGA outperforms NTAC in both accuracy and macro-F1 on all datasets, with particularly large gains in macro-F1 in the visual datasets. Transfer further improves accuracy in all four hemisphere pairs, with the largest improvements in the female datasets. In the central brain and ventral nerve cord, recovery remains less accurate than in the visual datasets.

In Fig. 6, we examine recovery by expert-type size. MOMONGA considerably improves recovery of small types in the visual datasets, showing that its gains are not confined to types containing many neurons. In the central brain and nerve cord, some size ranges show little improvement over NTAC, and substantial errors remain.

**Figure 6.**
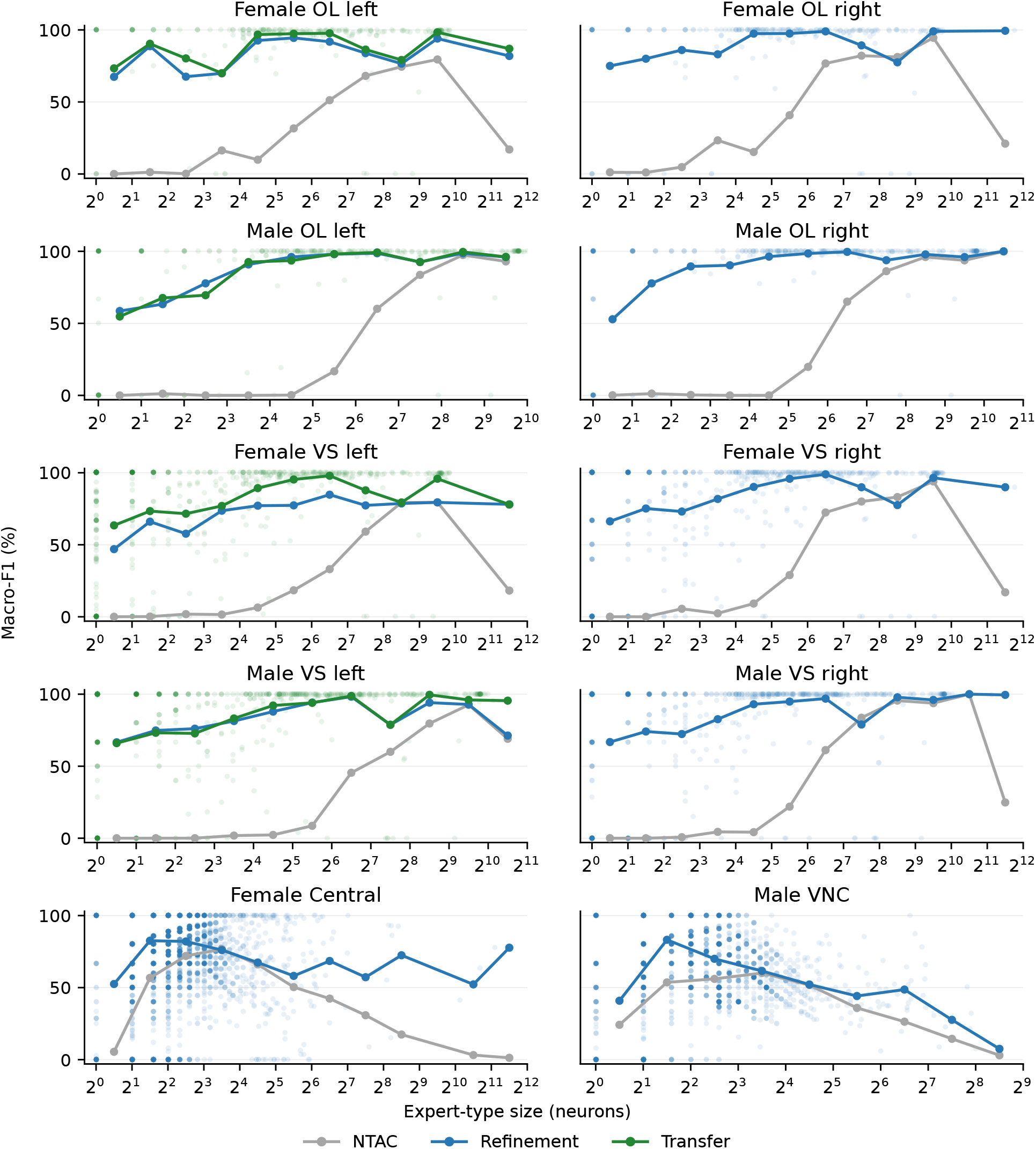
Recovery by expert-type size. F1 is computed for each expert type after one-to-one matching of the full partition to the expert annotations; unmatched types score zero. Curves show macro-F1 within size ranges containing 1, 2–3, 4–7, 8–15 neurons and successive doubling ranges, with equal weight per type. All curves use the selected partitions; no partition is selected separately for each type-size range. Dots correspond to individual types in the final MOMONGA output (green when transfer is applied). Curve points sit at the centre of their size range, on the log axis. NTAC and refinement are deterministic; transfer curves are means over five sets of random seeds.

### Transfer between specimens

To show that transfer also works across specimens, we use the male connectome as donor for the female connectome: Female OL left and Female VS left receive transfers from the Male left and Male right partitions of the same region (Fig. 7). Evaluation covers all connected recipient neurons, including types absent from the donor specimen. In every run the algorithm correctly chooses the male partition as the donor. Both male donors raise accuracy and macro-F1 well above refinement alone, matching or exceeding the female right hemisphere as donor. Transfer is therefore as effective between specimens of different sexes as between hemispheres of one specimen.

**Figure 7.**
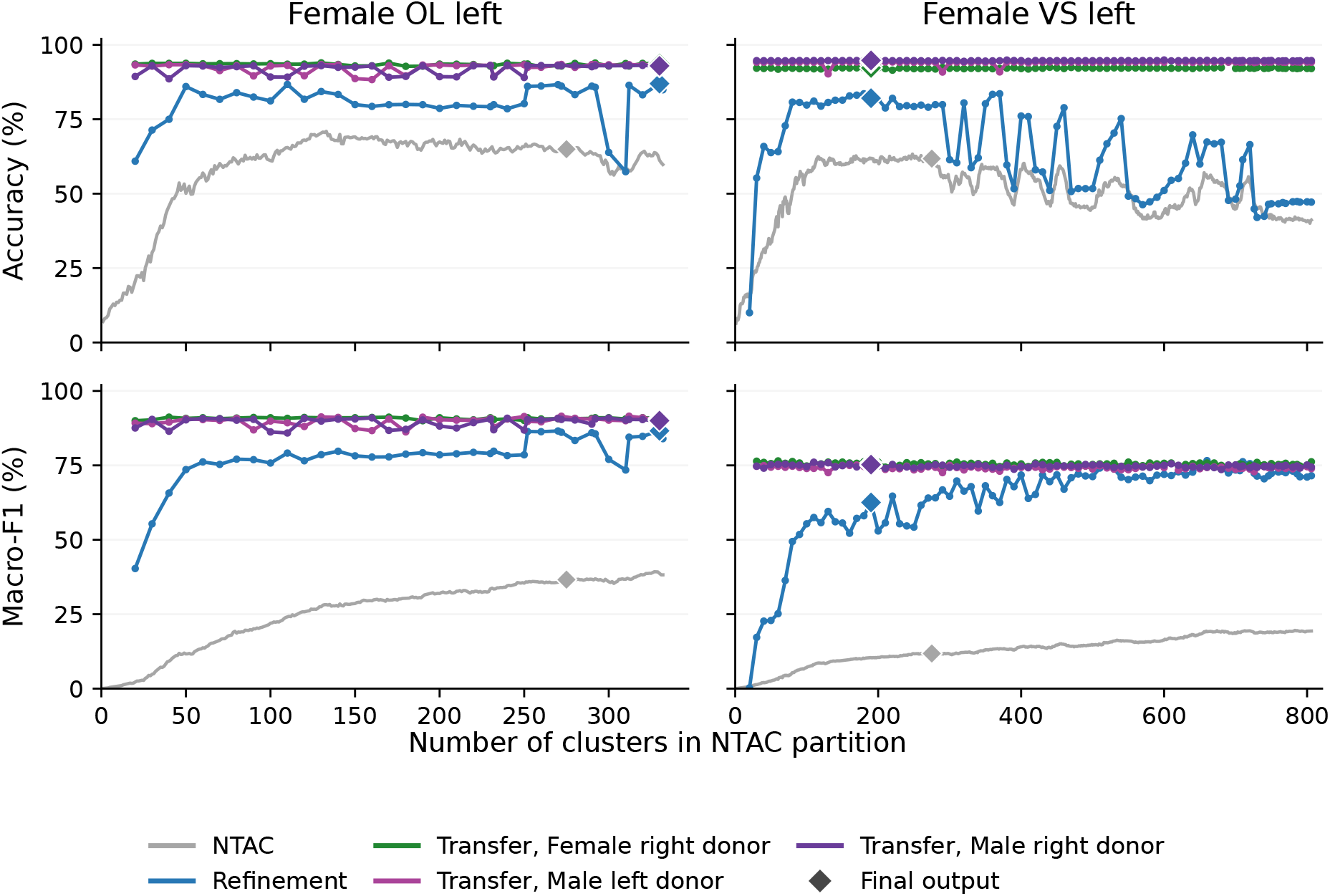
Transfer between specimens: Female OL left and Female VS left receive transfers from the Male left and Male right hemispheres of the same region, compared with NTAC, refinement alone and transfer from the female right hemisphere. Curves show accuracy and macro-F1 across NTAC starting partitions. Diamonds mark the final outputs: the minimum-Jaccard NTAC partition and the MDL-selected MOMONGA partition, after transfer where applicable. The donors are the final MOMONGA partitions of the male hemispheres (denoted by diamonds in Fig. 5). All connected recipient neurons are scored, including types absent from the male specimen. NTAC and refinement are deterministic; transfer diamonds are means over five sets.

## Discussion

Our findings indicate that MOMONGA is the first reliable unsupervised method for full cell typing in large regions of the fly brain. In regions where recovery is incomplete, the confidence score isolates a subset of neurons that forms pure representatives of many expert types, saving considerable human effort in the typing process. Transfer raises the weaker of two partitions to the level of the stronger without labels on either side, between hemispheres and between specimens of different sexes; we believe it can be a useful primitive beyond the scope of this work.

Our MDL framework is not tied to the particular code used here. The search, selection and transfer operate on any encoding of the connectome given a partition. Typing in a new region, or with additional data, requires only a different encoding, not a different method. This is the natural route towards improved typing of the central brain and nerve cord.

## Methods

To present MOMONGA, we first introduce notation, then describe the code that defines the description length, and finally the refinement and transfer algorithms.

### Notation and definitions

Let *G* = (*V, E, w*) be a directed weighted graph on *n* = |*V* | vertices. Assume that every directed edge (*u, v*) has an integer weight *w*(*u, v*) *≥ b*, where *b ≥* 1 is a given integer. Set *w*(*u, v*) = 0 when (*u, v*) *∉ E*. For sets *A, B ⊆ V*,

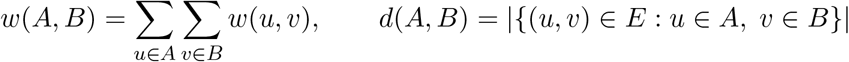

are the total edge weight and the number of edges from *A* to *B*; a vertex stands for its singleton, so *w*(*u, C*) is the total edge weight from *u* into *C* and *d*(*u, C*) is the number of vertices in *C* reached by *u*. The out- and in-weights are *w*^+^(*A*) = *w*(*A, V* ) and *w*^*−*^(*A*) = *w*(*V, A*). A partition *P* = {*C*_1_, …, *C*_*k*_} divides *V* into *k* clusters of sizes *n*_*i*_ = |*C*_*i*_|. We define

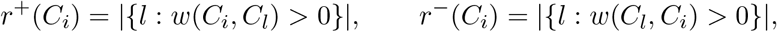

to be the numbers of clusters reached by *C*_*i*_ and reaching *C*_*i*_, respectively. Logarithms are in base 2 throughout.

### Jaccard distance

For two non-negative vectors *x* = (*x*_1_, …, *x*_*m*_) and *y* = (*y*_1_, …, *y*_*m*_), define their weighted Jaccard distance by

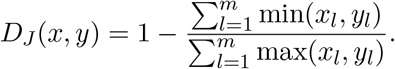

When both vectors are zero, we set *D*_*J*_ (*x, y*) = 1.

### Connectivity profiles

For a set of vertices *A ⊆ V*, its connectivity profile with respect to *P* is the non-negative vector *f*_*P*_ (*A*) of length 2*k*:

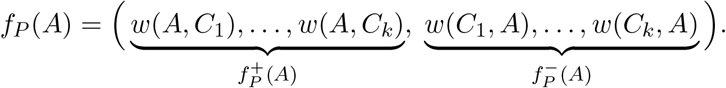

Its first *k* entries, 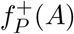, give the total outgoing edge weight from *A* to each cluster; its last *k* entries, 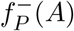, give the total incoming edge weight from each cluster to *A*. A vertex stands for its singleton: *f*_*P*_ (*u*) = *f*_*P*_ ({*u*}), and *f*_*P*_ (*A*) =Σ _*u∈A*_ *f*_*P*_ (*u*). Define the normalized profiles by

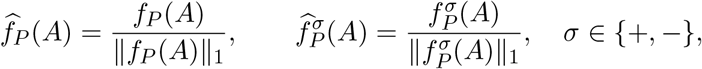

where ∥ *x* ∥ _1_ = Σ_*l*_ |*x*_*l*_| is the,*l*_1_ norm; when a denominator is zero, set the corresponding normalized profile to zero.

### The code

#### Setup

To define the code we assume two parties, a sender and a receiver. The sender knows the graph *G* = (*V, E, w*) and a partition *P* . The sender aims to use the partition to encode and transmit the graph *G* as efficiently as possible. For simplicity, we assume some shared information: *V* and the outgoing and incoming totals *w*^+^(*u*) and *w*^*−*^(*u*) for every *u ∈ V* are known to both parties.

Our code consists of four sequential stages. Roughly speaking, the sender first sends the partition, *P*, then the cluster-to-cluster connectivity, *w*(*C*_*i*_, *C*_*j*_) for all pairs of clusters, then for every vertex *u ∈ V* its weights to each cluster, *w*(*u, C*_*j*_), and finally the individual edges per vertex. Each stage uses the fact that the receiver already knows the information from the previous stage. The last three stages describe the graph from the outgoing side of its edges, by what each cluster and then each vertex sends. Describing it from the incoming side, by what each cluster and each vertex receives, gives a different code length, and we average the two. The graph *G* is fixed throughout this section, so every description length is presented as a function of the partition *P* .

#### The partition

To send the partition *P* = {*C*_1_, …, *C*_*k*_}, the sender first sends *k*, followed by the ordered cluster sizes *n*, …, *n* . These are positive integers summing to *n*, so there are 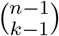 possible size lists. Given those sizes, the sender specifies which vertices belong to each cluster. There are *n*!*/*(*n*_1_! *· · · n*_*k*_!) assignments with the prescribed sizes. The cost in bits for this stage is

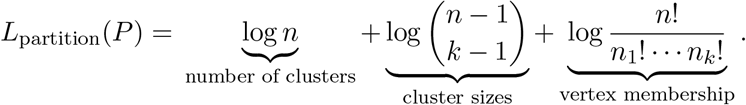

#### Cluster profiles

This stage describes how we send the total edge weight from each cluster to every cluster, including itself. For each *C*_*i*_, the sender describes its outgoing profile 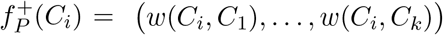.

The receiver already knows its total *w*^+^(*C*_*i*_), obtained by summing its members’ outgoing totals; this stage specifies how that total is divided among the *k* clusters. We use the cheaper of a dense and sparse encoding for this.

In the dense case, the sender specifies the complete vector of *k* non-negative weights whose sum is *w*^+^(*C*_i_ ). There are 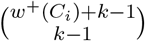 such vectors, giving the cost for *C* :

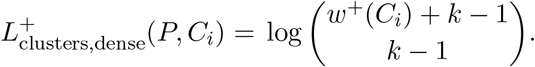

The dense description pays for all *k* coordinates, even when there are no edges to a certain cluster. For *k* = 1, the dense cost is zero.

In the sparse case the sender first gives *r*^+^(*C*_*i*_), the number of positive entries. It lies between one and min(*k, w*^+^(*C*_*i*_)). Next, the sender names their positions among the *k* entries, then specifies their weights, whose sum is *w*^+^(*C*_*i*_). These three steps give the cost for *C*_*i*_:

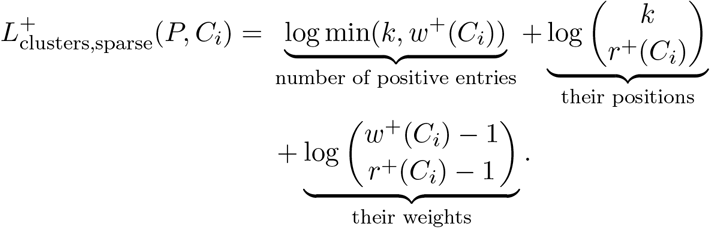

The sparse description pays only for positive entries: their number, positions and weights. The sender uses the cheaper encoding for each positive-total profile, with one bit specifying which. Zero-total profiles are already known to be zero. Thus

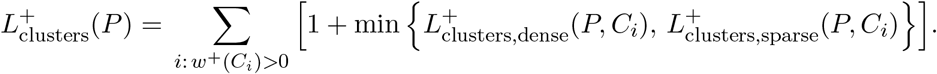

The incoming description 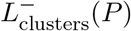 is defined analogously for incoming profiles, using *w*^*−*^(*C*_*i*_) and *r*^*−*^(*C*_*i*_) in place of *w*^+^(*C*_*i*_) and *r*^+^(*C*_*i*_).

#### Vertex profiles

For each cluster *C*_*i*_, the receiver already knows the total outgoing weight *w*^+^(*u*) of every vertex *u ∈ C*_*i*_, and from the preceding stage knows the total weight *w*(*C*_*i*_, *C*_*l*_) sent from *C*_*i*_ to every destination cluster *C*_*l*_. This stage describes how those cluster-level totals are divided among the individual vertices of *C*_*i*_.

Let 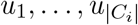 be an ordering of the vertices in *C*_*i*_ according to their indices. Form a list, *ℒ*, of length *w*^+^(*C*_*i*_): the first *w*^+^(*u*_1_) entries correspond to *u*_1_, the next *w*^+^(*u*_2_) entries to *u*_2_, and so on. Denote by *ℒ* (*u*) the sublist of *ℒ* corresponding to vertex *u*. An entry corresponding to *u* is labelled *l* when that unit of outgoing weight from *u* goes to cluster *C*_*l*_. Since the cluster-level totals are already known, the label *l* must occur exactly *w*(*C*_*i*_, *C*_*l*_) times in *ℒ*.

Choosing successively the positions carrying labels 1, 2, …, *k* gives the number of lists consistent with the cluster-level totals:

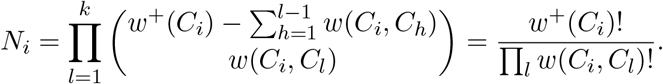

Expanding the binomial coefficients into factorials makes the product telescope: the factorial of the number of positions remaining after one choice cancels with the numerator factorial in the next choice.

The order of labels within *ℒ* (*u*) is irrelevant and only the counts *w*(*u, C*_*l*_) matter. The number of such orderings is

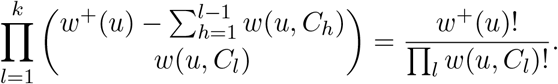

The same telescoping cancellation gives the factorial ratio here. Consequently, the over-counting factor for *N*_*i*_ is given by

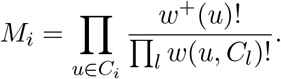

Since the order within each *L*(*u*) carries no information, we discount these *M*_*i*_ equivalent orderings. The resulting description length is log *N*_*i*_ *−* log *M*_*i*_ = log(*N*_*i*_*/M*_*i*_). Summing over clusters,

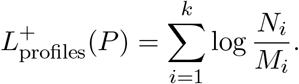

The incoming term 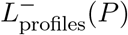 is defined symmetrically for incoming profiles.

#### Individual connections

For every vertex *u ∈ V* and cluster *C*_*l*_ *∈ P* with *w*(*u, C*_*l*_) *>* 0, the sender names how many vertices of *C*_*l*_ it reaches, which ones, and how its total weight *w*(*u, C*_*l*_) is divided over them. Recall that each edge has weight at least *b*. The number of possible partners of *u* in *C*_*l*_ is |*C*_*l*_ \ {*u*}|; we exclude *u* itself because the graph has no self-edges. There are at most min(|*C*_*l*_ \ {*u*}|, *lw*(*u, C*_*l*_)*/bj*) possible partner counts. Subtracting *b −* 1 from each of the *d*(*u, C*_*l*_) weights leaves positive integers with total *w*(*u, C*_*l*_) *−* (*b −* 1)*d*(*u, C*_*l*_), giving

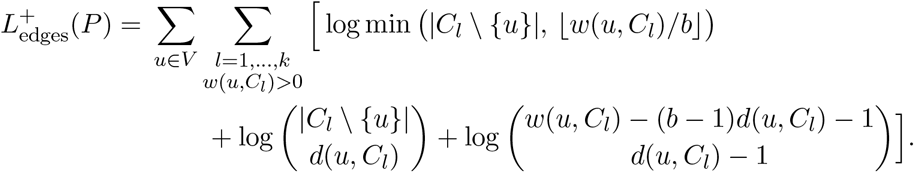

The incoming expression 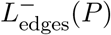 is defined symmetrically and uses source-cluster sizes.

#### Putting everything together

For the last three stages we average the outgoing and incoming descriptions, so that the code length does not depend on the choice of direction:

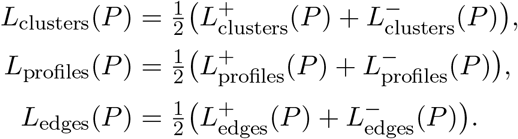

The total description length is

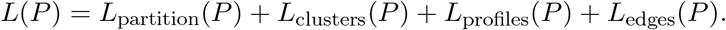

#### Interpretation

For probability vectors *p, q* of length *k*, define the Kullback–Leibler divergence by [21, Chapter 2]

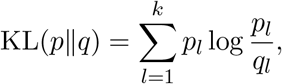

where terms with *p*_*l*_ = 0 contribute zero; if *p*_*l*_ *>* 0 and *q*_*l*_ = 0, the divergence is infinite. This divergence is nonnegative and vanishes exactly when *p* = *q*. The outgoing vertex-profile cost satisfies

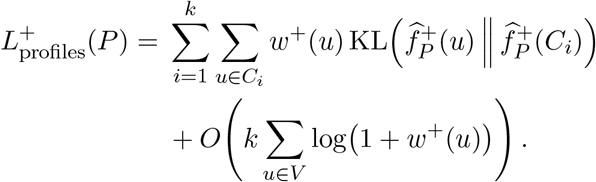

The incoming expression is symmetric, and *L*_profiles_(*P* ) averages the two. Its leading contribution therefore measures disagreement between normalized vertex profiles and their cluster profiles, weighted by outgoing and incoming totals. This contribution vanishes when every vertex’s normalized profile agrees with its cluster’s. The exact cost also includes corrections that can remain nonzero when the profiles agree. The full derivation and error bound are given in Appendix A.2.

## MOMONGA: refinement and transfer

### Refinement

The input to the refinement process is a graph *G* = (*V, E, w*), partition *P* = {*C*_1_, *C*_2_, …, *C*_*k*_} and an optional partition of the graph into left and right sides, *V*_left_, *V*_right_, such that *V*_left_ ∪ *V*_right_ = *V* and *V*_left_ *∩ V*_right_ = ∅.

Below we describe the building blocks of the algorithm, followed by a pseudocode description of the full refinement process in Algorithm 1.

#### Bilateral profiles

When left and right side labels are available we aim to use symmetry to get more informative profiles to be used in the next step. In the central brain and the nerve cord, *twin* neurons are common. A twin of a neuron *u* is a neuron *v* of the same type on the opposite side of the brain, with a connectivity pattern that mirrors that of *u*. That is, when neuron *u* connects to a neuron of type *A* on the left, its twin *v* connects to the corresponding neuron of type *A* on the right. The connectivity profiles of twins are therefore similar as long as the clusters of *P* are not split by side. But *P* may represent *A* as two one-sided clusters *A*_*L*_ and *A*_*R*_; then a neuron connected to *A*_*L*_ and its twin connected to *A*_*R*_ place the same connection in different coordinates, and their profiles look different although they should be grouped together. The construction below pairs vertices across sides, which is possible as long as *P* is not one-sided everywhere; applied to the members of *A*_*L*_ the pairing roughly recovers *A*_*R*_, and we use it to place the connections of twins in comparable coordinates.

We first construct the complete bipartite graph between *V*_left_ and *V*_right_, assigning edge (*x, y*) the weight *D*_*J*_ *(f*_*P*_ (*x*), *f*_*P*_ (*y*) ). Next, we run a greedy matching: repeatedly match pairs whose edge has minimum weight at both endpoints, then remove the matched vertices. Ties are resolved by vertex index; edges of weight one are excluded. Define *τ* : *V → V* by setting *τ* (*x*) = *y* and *τ* (*y*) = *x* for each matched pair *x, y*. For unmatched vertices, set *τ* (*v*) = *v*. Also define

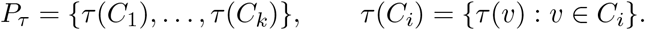

The *i*-th cluster of *P*_*τ*_ is *τ* (*C*_*i*_). The mapping *τ* maps each vertex to its likely twin on the opposite side, or to itself if none is found. The partition *P*_*τ*_ is a transformation of *P* with the clusters mapped under *τ* .

Using the above we can define the bilateral profile of a vertex. For each vertex concatenate the profiles under *P* and *P*_*τ*_, reversing their order between sides:

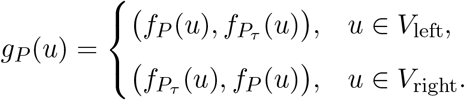

The resulting profile has 4*k* entries. The order is reversed so that the profiles of twins remain similar.

#### Nearest neighbours and shattering

Given a partition *P*, define the nearest neighbour of *u ∈ V* by

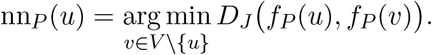

Ties are resolved by choosing the vertex with the smallest index. The nearest-neighbour relation need not be symmetric: it is possible that nn_*P*_ (*u*) = *v* but nn_*P*_ (*v*) *≠ u*.

When side information is available, we use *g*_*P*_ (*u*) instead of *f*_*P*_ (*u*) when finding nearest neighbours for shattering. The graph and *L*(*P* ) remain unchanged and *g*_*P*_ (*u*) is not used beyond the shattering step.

We define the *shattering* of a partition *P*, Shatter(*P* ), as the partition of *V* into connected components of the undirected graph with vertex set *V* and edge set {{*u*, nn_*P*_ (*u*)} : *u* ∈ *V* }. In other words, two vertices belong to the same cluster in the shattering of *P* if and only if a path of nearest-neighbour links connects them. Shattering produces a new partition by joining vertices connected through these links; its clusters may cross the boundaries of *P* .

#### Merging

The goal of merging is to combine clusters whose union has a shorter description length than the clusters separately. For distinct clusters *C*_*i*_, *C*_*j*_ *∈ P*, their merger gives the partition

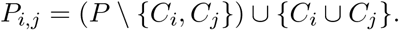

Ideally, we would apply the single best merger, recompute all pairs, and repeat; this is computationally expensive. Instead, merging proceeds in passes: each pass selects a set of candidate pairs, and before applying each merge we only verify that it still decreases *L*(*P* ) under the updated partition, without searching for the best one. We use the mutual-best rule as the heuristic for selecting candidates. At the beginning of each pass, each cluster chooses the partner whose merger gives the smallest *L*(*P*_*i,j*_), resolving ties in a fixed order. We retain pairs that choose each other and whose merger decreases *L*(*P* ), and process them in order of the decrease computed at the beginning of the pass. We then recompute the candidate pairs and repeat until no pairwise merger improves the description length. The resulting partition is Merge(*P* ).

#### Splitting

The goal of splitting is to separate a cluster *C*_*i*_ whose vertices have different connectivity profiles into two clusters. The exact change in *L*(*P* ) under a split is costly to compute, so we use a heuristic proposal based on Kullback–Leibler divergence. We define Split(*P* ) by splitting each cluster *C*_*i*_ *∈ P* into two clusters using the weighted KL 2-means algorithm [22].

During each split, the profile coordinates defined by *P* remain fixed. The algorithm uses repeated Lloyd-style iterations to partition *C*_*i*_ into two disjoint clusters, *A, B*, seeking to reduce the following objective:

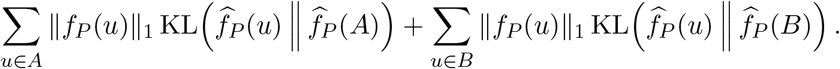

The two initial centres are the member furthest from *f* _*P*_ (*C*_*i*_) under KL divergence and the member furthest from that one; the iterations stop when the assignment is unchanged or after fifty. If either cluster is empty, *C*_*i*_ remains unchanged.

##### Algorithm 1

Refine(*G, P* )

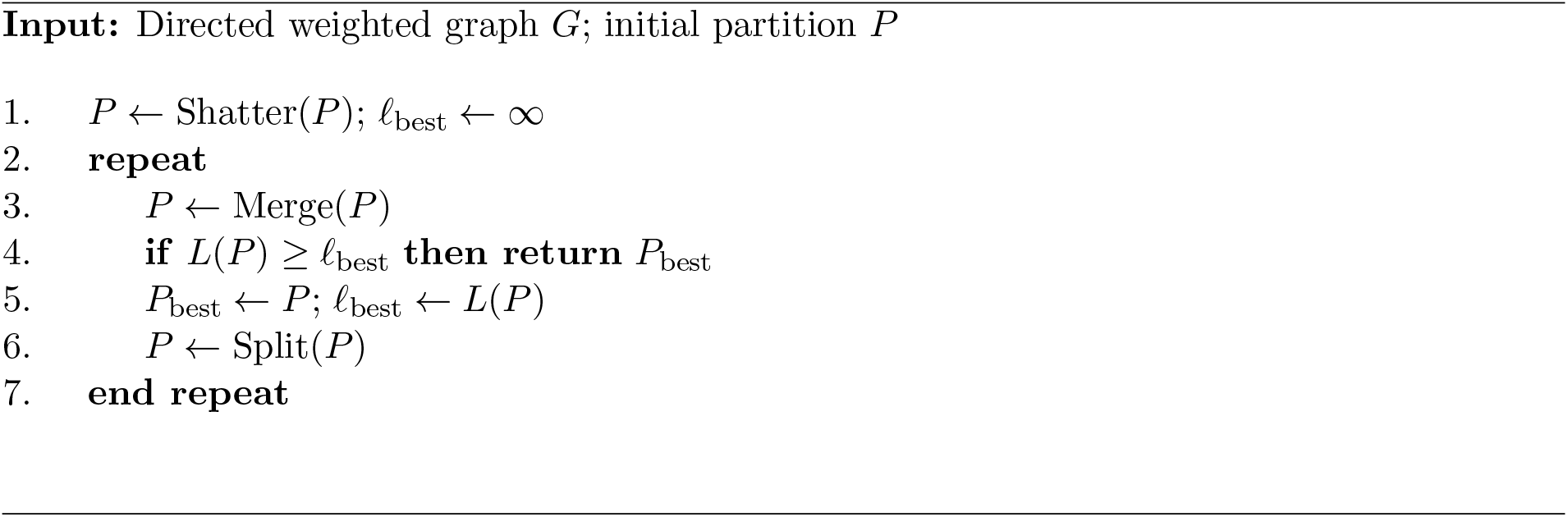

### Transfer

The input to the transfer procedure is two vertex-disjoint directed weighted graphs, *G*_1_ = (*V*_1_, *E*_1_, *w*_1_) and *G*_2_ = (*V*_2_, *E*_2_, *w*_2_), with partitions *P*_1_ and *P*_2_. We assume that one graph has a better partition than the other, and we want to use that partition to improve the other. The transfer process is unsupervised, and does not require any labels on either side. Until now we used *L*(*P* ) to measure the quality of a partition *P* on a fixed graph *G*. In this section we must explicitly state the graph the description length is evaluated on, and we write *L*(*P, G*) for the description length of partition *P* evaluated on graph *G*.

Below we describe the building blocks of the algorithm, followed by pseudocode in Algorithm 2.

#### Deciding the graph roles

Initially, we do not know which graph has the better partition. We use the description length to decide which graph is the donor and which is the recipient.

Define the description length per edge by *λ*_*t*_ = *L*(*P*_*t*_, *G*_*t*_)*/*|*E*_*t*_|, for *t ∈ {*1, 2}. The graph with the smaller *λ*_*t*_ is the *donor G*_*d*_, and the other is the *recipient G*_*r*_. Write *G*_*d*_ = (*V*_*d*_, *E*_*d*_, *w*_*d*_) and *G*_*r*_ = (*V*_*r*_, *E*_*r*_, *w*_*r*_). Let 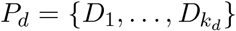 be the donor partition and 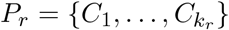 the recipient partition. And let DecideRoles(*G*_1_, *P*_1_, *G*_2_, *P*_2_) be the procedure that returns the donor and recipient graphs and partitions.

#### Initial matchings

We would like to compare profiles between the two graphs, but the coordinates of the profiles are defined by the clusters themselves, and we do not know which clusters correspond to each other. Our goal is to map every recipient cluster to a donor cluster with a similar connectivity profile. Several recipient clusters may map to the same donor cluster. If the correspondence between clusters were known, we could compare their profiles in common coordinates. However, those coordinates are themselves defined by the clusters we are trying to match.

For example, with two clusters in each graph, every recipient vertex has profile

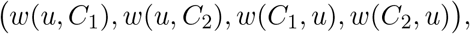

while every donor vertex has profile

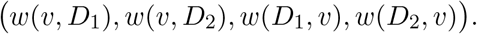

Both vectors have four entries, but we do not know whether the coordinates for *C*_1_ correspond to those for *D*_1_ or *D*_2_. We first obtain an initial correspondence from the connections between clusters, then use it to compare profiles and update the correspondence.

Let *A*^+^ be the *k*_*r*_ *× k*_*r*_ matrix whose *i*th row is 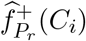, and let *A*^*−*^ be the matrix whose *j*th column is 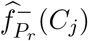):

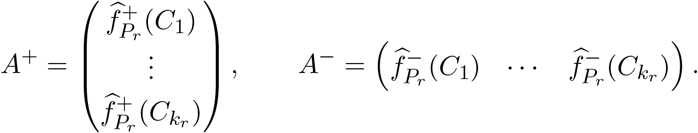

Entry (*i, j*) of both matrices describes the connection from *C*_*i*_ to *C*_*j*_: *A*^+^ is its fraction of the outgoing weight of *C*_*i*_, and 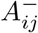 is its fraction of the incoming weight of *C*_*j*_. Define *B*^+^, *B*^*−*^ in the same way from the donor profiles. Finally, let *A* = *A*^+^ + *A*^*−*^ and *B* = *B*^+^ + *B*^*−*^, where we pad the smaller matrix with zero rows and columns such that both matrices have dimensions *k*_max_ *× k*_max_, where *k*_max_ = max(*k*_*r*_, *k*_*d*_).

We use the fast approximate quadratic assignment algorithm (FAQ) [23] to find a permutation *π* seeking to minimize

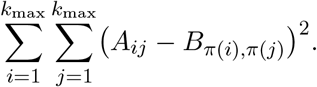

This objective compares connections between recipient clusters with connections between their proposed donor counterparts. FAQ returns a permutation of the padded cluster sets; discarding pairs that involve a padded cluster leaves exactly *k*_min_ = min(*k*_*r*_, *k*_*d*_) matched pairs. We relabel the clusters of both partitions so that matched pairs come first and share an index, giving the initial matching 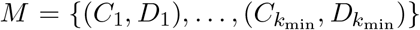, with unmatched clusters listed afterward. Relabelling changes only the order of the clusters, not their membership. We run FAQ, with at most sixty iterations per run, from *n*_init_ = 11 initializations, one uniform initialization and ten randomized ones, producing *n*_init_ candidate matchings, each with its own labelling. Let InitialMatchings(*G*_*r*_, *P*_*r*_, *G*_*d*_, *P*_*d*_, *n*_init_) be the procedure that returns these matchings.

#### Matching refinement

Given a matching *M*, we refine it using profile overlap. For a cluster *C* belonging to either partition *P*, define its aligned profile by

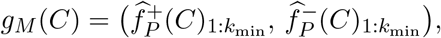

where 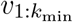 denotes the first *k*_min_ entries of a vector *v*. Both graphs now have 2*k*_min_-entry profiles in corresponding coordinates. The entries retain their original normalization; we do not renormalize after omitting unmatched coordinates.

For example, if the recipient has three clusters and the donor has two, with matching *M* = *{*(*C*_1_, *D*_1_), (*C*_2_, *D*_2_)}, the aligned profiles use connections to and from these two matched pairs. Although *C*_3_ is unmatched, its aligned profile still describes its connections to and from *C*_1_ and *C*_2_, so it can be compared with either donor cluster. Connections involving *C*_3_ as a profile coordinate are omitted because that coordinate has no matched donor counterpart.

Construct the complete bipartite graph between *P*_*r*_ and *P*_*d*_, with edge weights

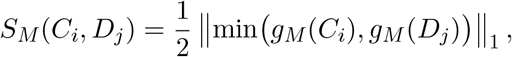

where the minimum is taken coordinatewise. *S*_*M*_ (*C*_*i*_, *D*_*j*_) is the fraction of weight the two aligned profiles share, averaged over the outgoing and incoming directions; it lies in [0, 1]. Find a maximum-weight matching *M*^*t*^ on this bipartite graph, which matches every cluster of the smaller partition. If *M*^*t*^ equals *M* or an earlier matching of this loop, stop and retain *M* . Otherwise, replace *M* by *M*^*t*^, reorder the clusters according to the new matching, recompute the scores, and repeat. Let FinalAlignment(*G*_*r*_, *P*_*r*_, *G*_*d*_, *P*_*d*_, *M* ) be the procedure that returns the final matching after this refinement.

#### Reassignment

Given a matching *M*, we label each recipient cluster with a donor cluster index, then move recipient vertices between the resulting *k*_*d*_ clusters according to profile similarity. The donor partition is fixed, and the recipient partition is updated.

First we extend the matching to a mapping *ϕ* : *{*1, …, *k*_*r*_*} → {*1, …, *k*_*d*_} by

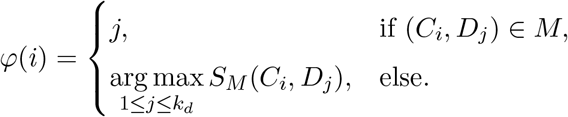

Define the recipient clusters

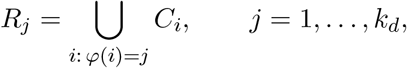

allowing empty clusters. In the three-to-two example, assigning *C*_3_ to donor index 1 gives *R*_1_ = *C*_1_ *∪ C*_3_ and *R*_2_ = *C*_2_. Write 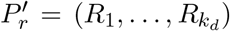 this recipient partition, retaining empty clusters; it is the starting point of reassignment, and *R*_*j*_ denotes its current *j*th cluster throughout. Recipient and donor profiles then have the same 2*k*_*d*_ coordinates, outgoing and incoming connections to corresponding clusters, with *R*_*j*_ corresponding to *D*_*j*_. The coordinates of an empty recipient cluster are zero.

We are now ready to reassign vertices between clusters. Define the mean cluster profiles

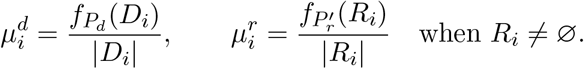

A donor update assigns each recipient vertex *u ∈ V*_*r*_ to a cluster minimizing 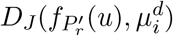. A joint update instead uses

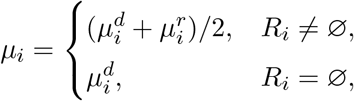

and assigns *u* to a cluster minimizing 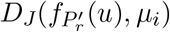. Each update assigns all recipient vertices simultaneously, using profiles and means computed from the partition before that update. The donor partition and its mean profiles remain fixed.

A *pass* consists of (1) one donor update, followed by (2) joint updates, repeated until the partition is unchanged or repeats one produced earlier in the pass; the pass ends with that partition. Starting from 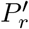, perform passes until a pass ends with the partition it started from, or with one that an earlier pass ended with, counting 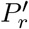 as an earlier result. Let Reassign(*G*_*r*_, *P*_*r*_, *G*_*d*_, *P*_*d*_, *M* ) be the procedure that returns this partition.

#### Selection and local search

Reassignment returns a recipient partition 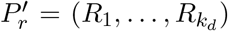 whose clusters are indexed by donor clusters, so it can be joined with *P*_*d*_ cluster by cluster.

Form the disjoint union *G*_*d*_ *G*_*r*_ and its joint partition *Q* = *{D*_*j*_ *∪R*_*j*_ : *j* = 1, …, *k*_*d*_*}. Q* has *k*_*d*_ clusters, each consisting of one donor cluster and its recipient counterpart; the union graph has no edges between the two graphs. Evaluate *L*(*Q, G*_*d*_ ⨆ *G*_*r*_) for each of the *n*_init_ candidates after reassignment, and select the candidate, *Q*_best_, with the smallest *L*(*Q, G*_*d*_ ⨆ *G*_*r*_).

Next, we improve the selected candidate, *Q*_best_, by local search on description length, while keeping the donor vertices in their original clusters and the *k*_*d*_ cluster indices fixed. At the start of each sweep, identify recipient vertices with an improving move. Visit these vertices in increasing order of index, recomputing each vertex’s best destination under the current partition and applying its move if it strictly decreases *L*(*Q, G*_*d*_ ⨆ *G*_*r*_). Repeat until no recipient vertex has an improving move. Let LocalSearch(*G, Q, V* ^*t*^) denote this procedure: starting from a partition *Q* of *G*, it moves only vertices of *V* ^*t*^ and returns the final partition. The recipient part of its output, *{A ∩ V*_*r*_ : *A ∈ Q}*, is the transferred partition.

##### Algorithm 2

Transfer(*G*_1_, *P*_1_, *G*_2_, *P*_2_)

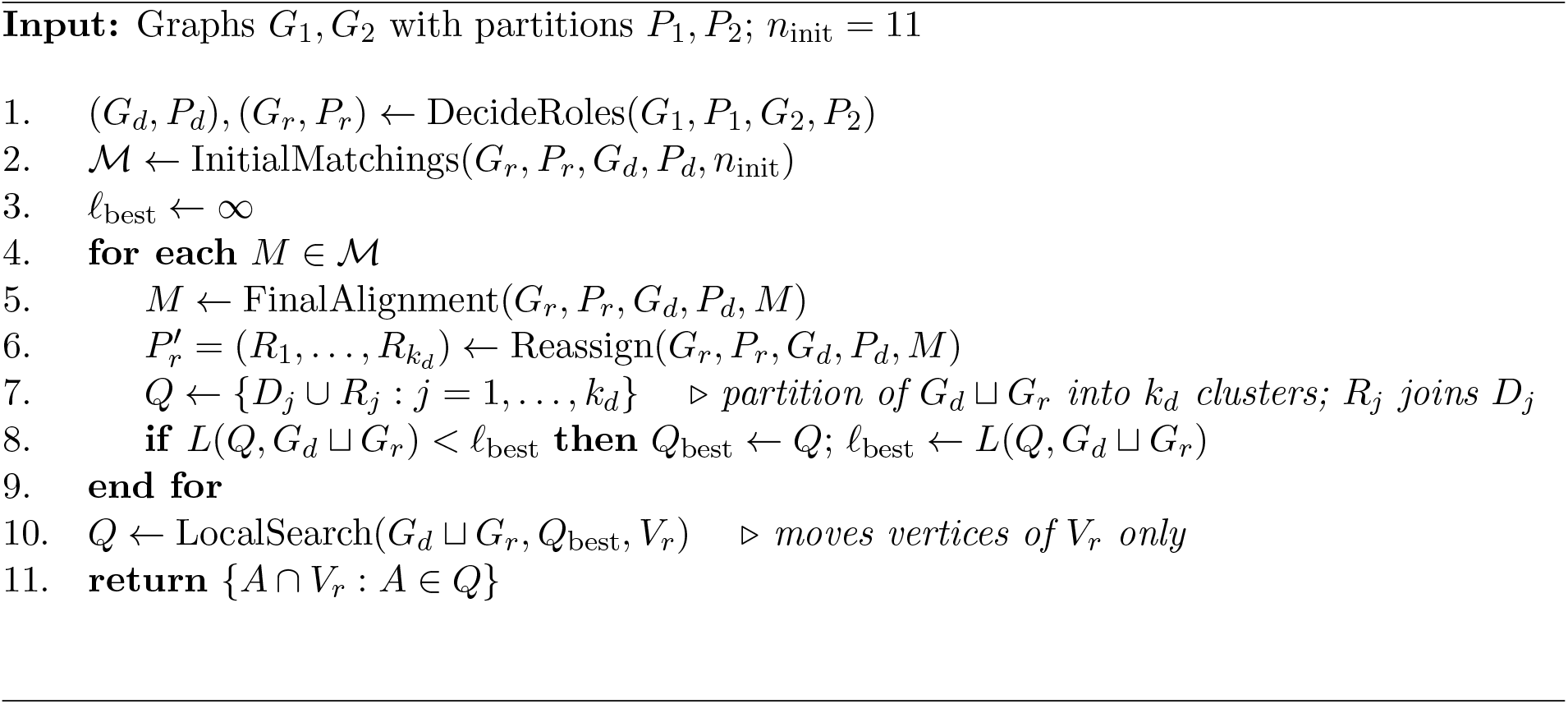

## Datasets

Dataset names combine the specimen’s sex (Female or Male) with its region, OL (optic lobe), VS (visual system), Central (central brain) or VNC (ventral nerve cord), and the hemisphere, left or right, where hemispheres are analysed separately. Female OL, Female VS and Female Central use the FAFB reconstruction with FlyWire segmentation v783, downloaded from Codex (https://codex.flywire.ai); expert types are the primary types of its consolidated cell-type table, and region membership and side come from the NTAC release [11]. Male OL and Male VS use the per-side MaleCNS exports from flywire.ai; expert types are their cluster annotations. Male VNC uses the MANC neuron and connection tables; expert types are its primary cell types, and side is the soma side. Every graph retains only connections of weight at least *b* = 5, the threshold of the published FlyWire network analysis [1]. Accuracy and macro-F1 are computed over the neurons with at least one retained connection; Supplementary Table 1 lists these populations.

### Statistics and reproducibility

All analyses used publicly available reconstructions of Drosophila connectomes (central brain, optic lobes and ventral nerve cord). Sample sizes were not predetermined; they reflect the full datasets or anatomically defined subsets described in the Methods. No data was excluded. Because this study uses only fixed connectivity datasets, standard biological randomization and blinding do not apply. For transfer experiments performance metrics (accuracy and macro-F1) were averaged over five runs with different random seeds.

## Data and code availability

The connectome datasets are publicly available: the FAFB female brain at https://flywire.ai and https://codex.flywire.ai, the male central nervous system at https://male-cns.janelia.org, and MANC at https://www.janelia.org/project-team/flyem/manc-connectome. MOMONGA is available as a Python package at https://github.com/gregoryschwartzman/momonga, together with the graphs analysed here. The package also contains a much faster implementation of NTAC that produces the same partitions as the original.

## Acknowledgements

The author was supported by KAKENHI 25K00370, JST ASPIRE JPMJAP2302 and JST CRONOS JPMJCS24K2.

## A Omitted proofs and definitions

### A.1 Confidence

For a partition *P* = {*C*_1_, …, *C*_*k*_} and a vertex *u ∈ C*_*i*_, the confidence of *u* is

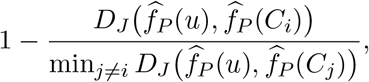

truncated to [0, 1]; a zero denominator gives confidence zero. It measures the margin between a vertex’s own cluster and the nearest other cluster, and uses no expert labels. The catalogues in Fig. 2 retain neurons in decreasing order of confidence, computed once on the fixed output partition, with the same score for MOMONGA and NTAC.

### A.2 Profile interpretation

We derive the relation between the vertex-profile description length and agreement between normalized vertex and cluster profiles, keeping an explicit bound on the approximation error. For a probability vector *p* = (*p*_1_, …, *p*_*k*_), define its entropy by

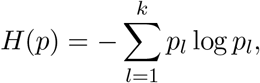

with 0 log 0 = 0. The following standard inequality relates entropy to the factorial ratios in the profile code.

#### Lemma A.1

(Multinomial coefficient bound). *For nonnegative integers x*_1_, …, *x*_*k*_ *with total* 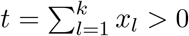. *Then [24, Lemma 2*.*3]*

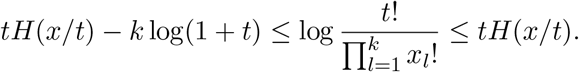

Applying this inequality to the vertex and cluster profiles gives the following bound on the difference between the exact cost and its KL expression.

#### Lemma A.2

(Vertex-profile cost). *The outgoing vertex-profile cost satisfies*

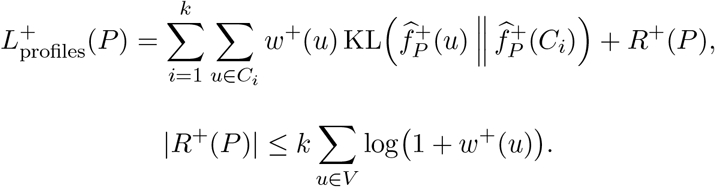

*Proof*. For a nonnegative integer vector *x* of length *k* and total *t* = *∥ x ∥*_1_ *>* 0, let

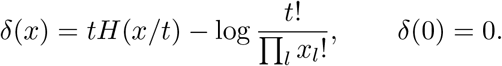

We have

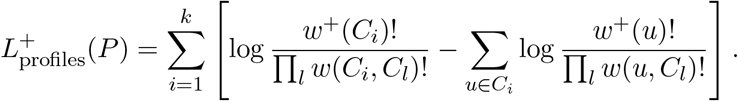

Fix a cluster *C*_*i*_ and consider its summand. Write

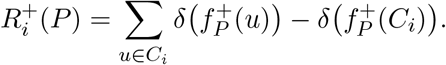

Then

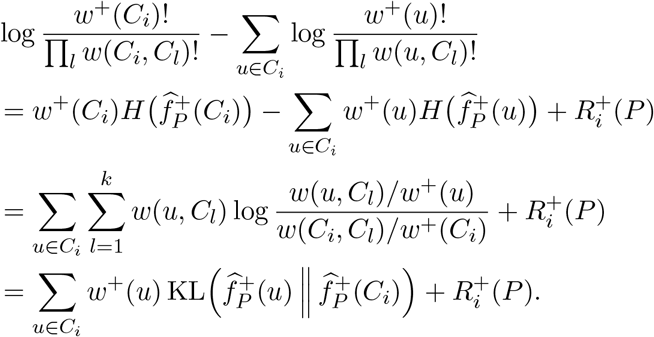

The first equality uses the definition of *δ* with 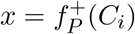 and 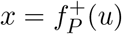. The second expands the entropies and uses 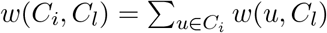. The third uses the definitions of normalized profiles and KL divergence.

Summing over clusters gives the claimed expression, with

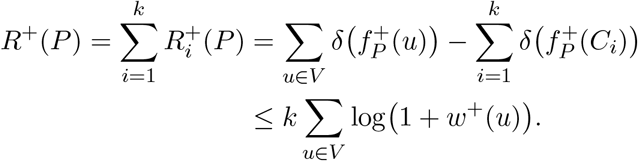

Lemma A.1 gives 0 *δ*(*x*) *k* log(1 + *t*). Both deficit sums lie between zero and *k* log(1 + *w*^+^(*u*)), since 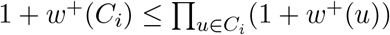. Their difference obeys the claimed bound.

The lemma holds symmetrically for incoming profiles, with + replaced by *−* throughout.

## Supplementary Material

**Supplementary Table 1.** Dataset populations after retaining connections of weight at least five: vertices, vertices left without a connection, and expert types among the rest.

| Dataset | Vertices | Isolated | Types |
| --- | --- | --- | --- |
| Female OL right | 43,175 | 9 | 232 |
| Female OL left | 43,430 | 622 | 231 |
| Female VS right | 47,404 | 25 | 724 |
| Female VS left | 47,600 | 632 | 705 |
| Female Central | 41,714 | 892 | 7,928 |
| Male OL right | 44,768 | 0 | 263 |
| Male OL left | 44,567 | 0 | 264 |
| Male VS right | 53,277 | 0 | 723 |
| Male VS left | 51,777 | 0 | 717 |
| Male VNC | 23,641 | 98 | 4,064 |

## Footnotes

1 NTAC takes as input a parameter *k* and iteratively creates partitions into 1, 2, 3, …, *k* clusters. We consider a subset of these partitions as inputs to MOMONGA: every 10th (every 100th for Female Central and Male VNC). The x-axis in Figs. 5 and 7 is the number of clusters in the input partition.

